# *C. elegans* LET-381/FoxF and DMD-4/DMRT control development of the mesodermal HMC endothelial cell

**DOI:** 10.1101/2024.12.23.630164

**Authors:** Nikolaos Stefanakis, Jasmine Xi, Jessica Jiang, Shai Shaham

## Abstract

Endothelial cells form the inner layer of blood vessels and play key roles in circulatory system development and function. A variety of endothelial cell types have been described through gene expression and transcriptome studies; nonetheless, the transcriptional programs that specify endothelial cell fate and maintenance are not well understood. To uncover such regulatory programs, we studied the *C. elegans* Head Mesodermal Cell (HMC), a non-contractile mesodermal cell bearing molecular and functional similarities to vertebrate endothelial cells. Here, we demonstrate that a Forkhead transcription factor, LET-381/FoxF, is required for HMC fate specification and maintenance of HMC gene expression. DMD-4, a Dmrt transcription factor, acts downstream of and in conjunction with LET-381 to mediate HMC fate specification and gene expression maintenance. DMD-4, independently of LET-381, also represses the expression of genes associated with a different, non-HMC, mesodermal fate. Our studies uncover essential roles for FoxF transcriptional regulators in endothelial cell development, and suggest that the identity of FoxF co-functioning target transcription factors promotes specific non-contractile mesodermal fates.

## INTRODUCTION

Endothelial cells are mesodermal cells that line blood vessels. They are crucial for maintaining circulatory system integrity, regulating blood flow, and maintaining ion homeostasis of surrounding tissues. Specialized capillary endothelial cells are key components of the vertebrate blood-brain barrier. Aberrant endothelial cell development and function is associated with various diseases including cancer, atherosclerosis and stroke (Rohlenova et al., 2018).

Endothelial cells originate from the embryonic mesoderm germ layer. During vasculogenesis, mesodermal precursors of both endothelial and hematopoietic cells aggregate into cell clumps called blood islands. Later, the outer layer of these clumps differentiates into endothelial cells, while inner cells give rise to hematopoietic cells. During subsequent angiogenesis, the endothelial cell population expands and remodels, sprouting and branching to form the vascular lining (Marziano et al., 2021). Although all vasculature beds exhibit common features, endothelial cells are phenotypically heterogeneous, functionally tailored to the specific needs of the tissue in which they reside (Hennigs et al., 2021). A large number of studies has defined key roles for vascular endothelial growth factor (VEGF) and Notch signaling in the early steps of vascular development (Olsson et al., 2006, Phng and Gerhardt, 2009). Furthermore, recent advances in multiomics technologies have facilitated molecular characterization of phenotypically-distinct endothelial cells, highlighting differences in gene expression and defining a host of endothelial subtypes (Trimm and Red-Horse, 2023). Nonetheless, the transcriptional programs involved in endothelial cell differentiation and subtype specification and maintenance remain largely unknown.

The nematode *C. elegans* has played pivotal roles in the identification of conserved molecular mechanisms of development and function of different tissues and cell types (Ambros, 2011, Gieseler et al., 2017, Horowitz and Shaham, 2024, Kratsios and Hobert, 2024, Richmond, 2005, Singhvi et al., 2024). The *C. elegans* Head Mesodermal Cell (HMC) was recently shown to bear similarities to vertebrate endothelia (Choi et al., 2023). Like endothelial cells, the HMC is a non-contractile, mesodermal cell. It resides within the animal’s main body cavity, the pseudocoelom, a simple circulatory system that provides a means for nutrient, oxygen, and signaling molecule distribution (Hall and Altun, 2008). G-protein-coupled receptors (GPCR) on the HMC respond to peptidergic signals to elicit changes in HMC intracellular calcium levels, leading to contraction of surrounding muscles connected to the HMC via gap junctions (Choi et al., 2023). Similarly, vertebrate endothelial cells can change their intracellular calcium levels in response to extracellular peptides and control smooth muscle contraction via myoendothelial gap junctions (Figueroa and Duling, 2009, Griffith, 2004, Maguire and Davenport, 2005).

To gain insights into the molecular mechanisms controlling specification and maintenance of endothelial cell fates, we therefore investigated HMC development. Anatomically, the HMC cell body lies dorsomedially, just above the posterior pharyngeal bulb (Fig. 1A). It extends a short anterior and a long dorsal posterior process as well as two lateral processes that project around the pharynx and merge ventrally, where they extend anteriorly and posteriorly to mirror the dorsal anterior/posterior processes. We found that LET-381, a FoxF transcription factor continuously expressed in the HMC, is required for both the initial specification and subsequent maintenance of HMC identity. The LET-381 target DMD-4, a DMRT transcription factor, acts with LET-381 to specify HMC fate and to maintain HMC gene expression. DMD-4 also represses expression of a gene normally expressed in GLR glia, a different mesodermal cell type. Previous studies showed that LET-381/FoxF acts with UNC-30/Pitx2 and CEH-34/Six2 to control the development of GLR glia and mesodermally-derived coelomocytes, respectively. Taken together, our studies reveal that LET-381/FoxF acts as a terminal selector factor for endothelial cell differentiation, and demonstrate that its co-functional target transcription factors specify the identities, and control maintenance, of distinct mesodermal fates.

**FIGURE 1.**
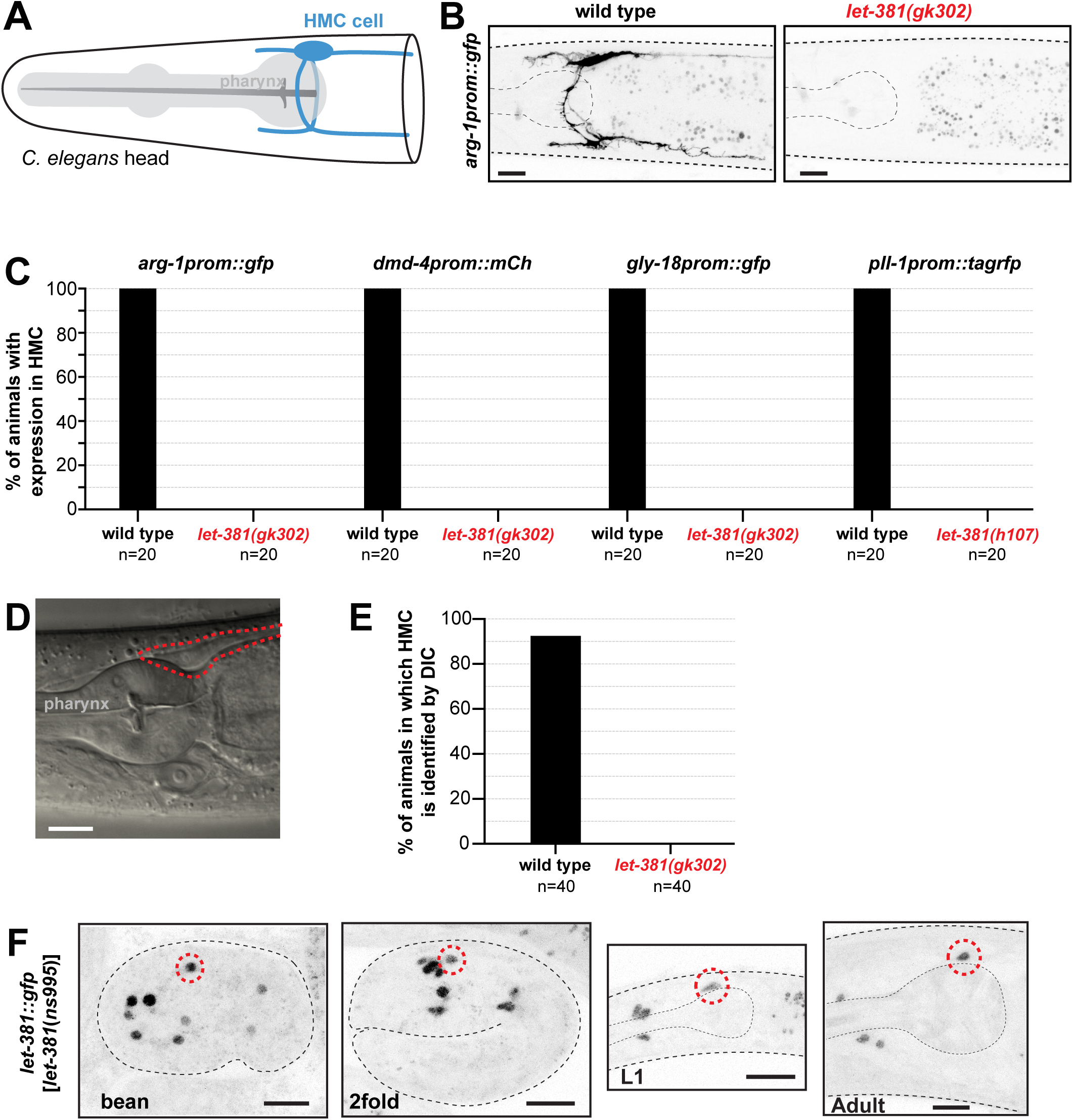
*let-381/Foxf* is required for HMC specification. (A) Schematic representation of the HMC cell (blue). (B) Fluorescence images of the HMC specific *arg-1prom::gfp* reporter in wild type (left) and *let-381*(*gk302*) mutant (right). Expression is not observed in the mutant. (C) Quantification of expression of four different HMC reporters (indicated above the bars) in wild type and *let-381* mutants. Expression in the HMC is not detected in the mutants. (D) Differential Interference Contrast (DIC) image of young adult wild-type animal posterior head region (see Fig. 1A). Red, The HMC cell body. (E) Percent young adult animals in which the HMC cell body can be identified by DIC microscopy in wild type and *let-381(gk302)* mutants. (F) Expression of *let-381::gfp* in embryonic stages (bean, 2-fold), L1 larva, and adult animals. Red, HMC. n=20 animals for each genotype in (C) and n=40 for each genotype in (E). Anterior is left, dorsal is up, and scale bars are 10 μm for all animal images.

## RESULTS

### *let-381/FoxF* is required for HMC cell specification

We previously reported that the LET-381/FoxF transcriptional regulator is expressed in GLR glia, coelomocytes, and the head mesodermal cell (HMC) (Amin et al., 2010, Stefanakis et al., 2024). While roles for LET-381 in GLR and coelomocyte development are well established (Amin et al., 2010, Stefanakis et al., 2024), whether LET-381 also promotes HMC specification was unknown. To address this question, we introduced HMC-specific reporter transgenes (*arg-1prom::gfp*, *dmd-4prom::mCherry*, *gly- 18prom::gfp*, *pll-1prom1::rfp*) into animals homozygous for the *let-381(gk302)* or *let- 381(h107)* alleles*. let-381(gk302)* animals contain a deletion removing LET-381 DNA binding domain encoding sequences; and *let-381(h107)* animals harbor a splice acceptor point mutation predicted to result in a truncated LET-381 protein (Barstead et al., 2012, Howell et al., 1987). While animals homozygous for *gk302* and *h107* undergo late-embryonic or early-larval developmental arrest, respectively, some escapers develop further to become sterile adults. We found that neither *let-381(h107)* arrested larvae nor *let-381(gk302)* adult escapers express any of the HMC reporters we tested (Fig. 1B, C). Furthermore, while the HMC cell body is easily identified by its characteristic shape and position in young adult wild-type animals using Differential Interference Contrast (DIC) microscopy, no HMC cell body is discerned in young adult escaper *let-381(gk302)* mutants (Fig. 1D, E). Taken together, these experiments suggest that LET-381 is required for HMC specification.

### *let-381/Foxf* is continuously and cell-autonomously required to maintain the HMC identity

To determine when and where LET-381 is required to promote HMC specification, we tracked expression of a CRISPR/Cas9-generated *let-381::gfp* reporter in which *gfp* coding sequences are inserted into the endogenous *let-381* locus. As shown in Figure 1F, *let- 381::gfp* expression in the HMC cell begins in comma-stage embryos and is maintained throughout adulthood, suggesting that LET-381 may be required not only to specify the HMC, but also to maintain its fate.

To test this idea, we sought to directly determine whether LET-381 is required for HMC fate maintenance. We previously showed that LET-381 promotes its own expression in GLR glia using an autoregulatory *let-381* binding motif located upstream of the first exon. A deletion/insertion genomic lesion removing this motif, *let-381(ns1026)*, does not affect LET-381 embryonic GLR glia expression, but blocks continued expression in these cells beyond the first larval stage (Stefanakis et al., 2024). While the *let- 381(ns1026)* mutation does not affect LET-381 expression in the HMC, another allele we generated, *let-381(ns1023)*, lacking sequences surrounding the autoregulatory *let-381* binding motif, preserves embryonic expression but blocks post-embryonic expression of LET-381 in both the HMC and GLR glia (Fig. 2A-C). Importantly, we found that expression of two endogenous and four transgenic reporters (*pll-1::gfp*, *gbb-2::gfp*, *snf-11^fosmid^::mCherry*, *gly-18prom::gfp*, *hot-2prom::gfp*, *glb-26prom::gfp*) is nearly entirely abolished in the HMC of *let-381(ns1023)* autoregulatory mutants (Fig. 3A, B). Expression levels of a *dmd-4prom::mCherry* HMC reporter are likewise substantially reduced (Fig. 3C, D). By contrast, expression of these reporters in the HMC is not affected in the GLR-specific *let-381(ns1026)* autoregulatory mutant. These observations support the conclusion that *let-381* acts cell autonomously to maintain HMC gene expression (Fig. 3A, B).

**FIGURE 2.**
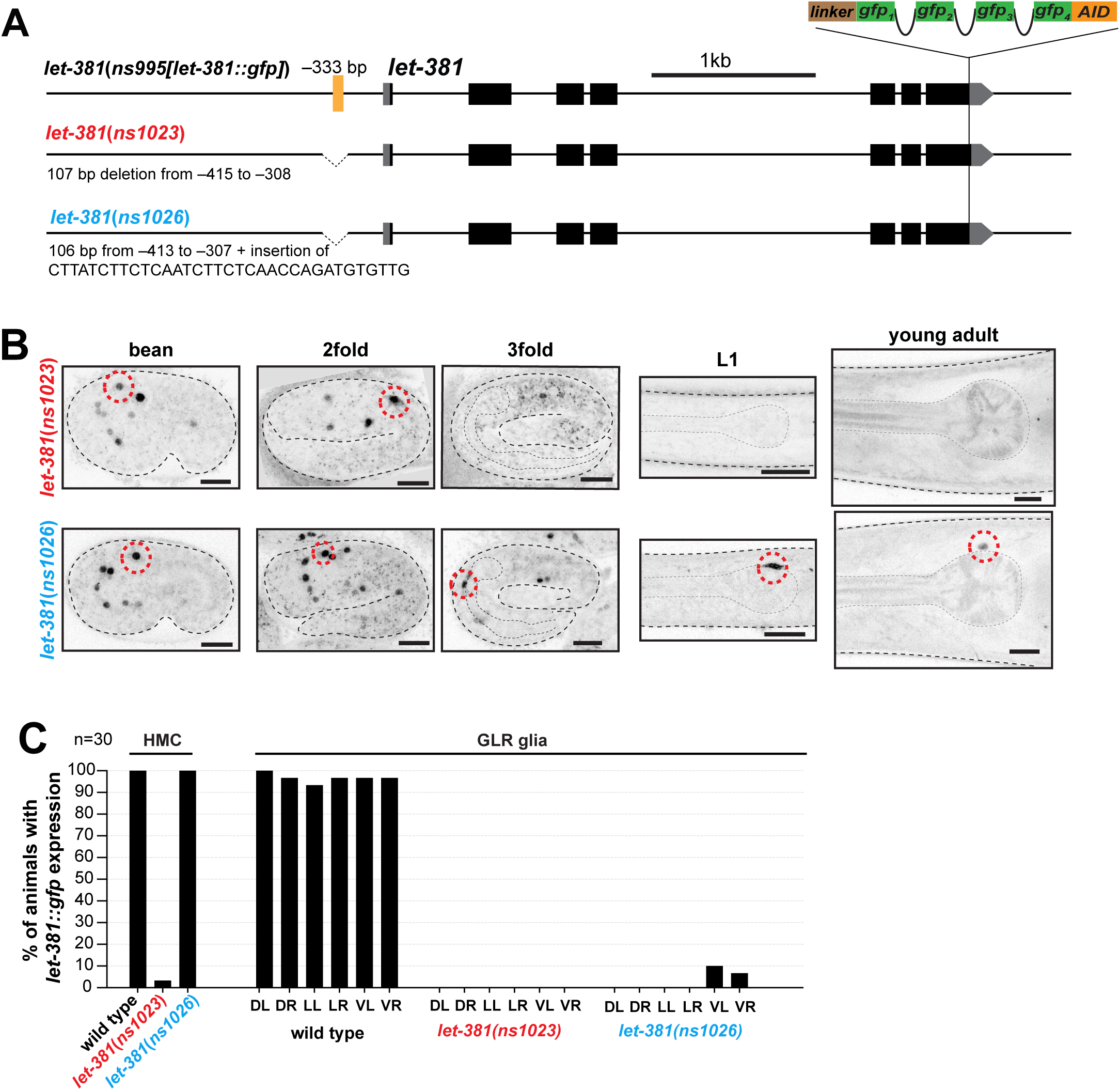
A *let*-*381* autoregulatory motif is required to maintain LET-381 expression in the HMC. (A) Schematics showing details of *let-381* autoregulatory motif deletion mutant alleles. Both alleles remove a sequence containing the *let-381* autoregulatory motif (yellow line), located at -333 bp from the ATG, but *let-381(ns1026)* also has an insertion. (B) Fluorescence images of *let-381::gfp* shown for bean, 2-fold and 3-fold embryonic stages, L1 larva, and young adults for each genotype. The *let*-*381*(*ns1023*) allele (upper panel row) affects maintenance of *let-381::gfp* expression in both HMC and GLR glia. *let*- *381*(*ns1026*) (lower panel row) affects maintenance only in GLR glia. Red, HMC. (C) Percentage of young adult wild-type, *let*-*381*(*ns1023*) and *let*-*381*(*ns1026*) animals with *let*-*381*::*gfp* expression in HMC and GLR glia. *let*-*381*::*gfp* expression is nearly abolished from both the HMC and GLR glia in *let*-*381*(*ns1023*). n=30 animals for each genotype scored in (C). Anterior is left, dorsal is up, and scale bars are 10 μm for all animal images.

**FIGURE 3.**
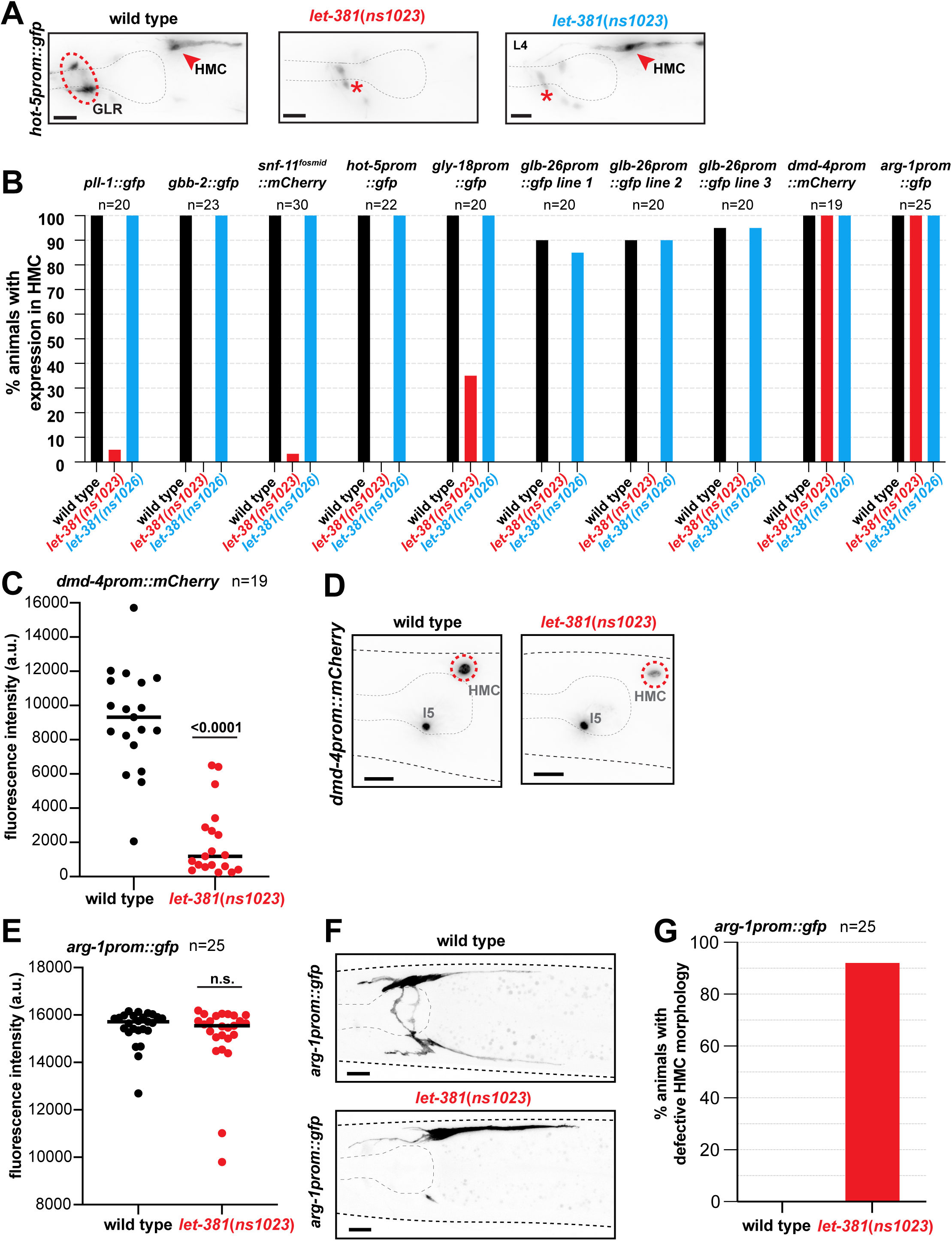
LET-381 is required for maintenance of HMC expression and morphology. (A) *hot*-*5*promoter::*gfp* is expressed in both HMC (red arrowhead) and GLR glia (red dashed circle) in wild-type animals. Expression of this reporter is lost in both cell types in *let*-*381*(*ns1023*) animals. By contrast, expression is lost only GLR glia in *let*-*381*(*ns1026*) animals. Red asterisk, unrelated expression of *hot*-*5prom*::*gfp* in neurons. (B) Percentage of animals with expression in HMC of eight different reporters (indicated above the bars) for wild type, *let*-*381*(*ns1023*), and *let*-*381*(*ns1026*). *glb-26*promoter::gfp is an extrachromosomal reporter and therefore three independent lines were tested. (C) Quantification of fluorescence intensity of *dmd*-*4*promoter::*mCherry* reporter in HMC in wild-type and *let*-*381*(*ns1023*) animals. *dmd*-*4*promoter::*mCherry* expression is significantly reduced in *let*-*381*(*ns1023*). Representative images for each genotype shown in (D). Red, HMC. (E) Quantification of fluorescence intensity of the *arg- 1*promoter*::gfp* reporter in wild-type and *ns1023* animals. Fluorescence intensity in the cell body is not affected in *let*-*381*(*ns1023*). Representative images for each genotype shown in (F). Lateral and ventral HMC processes are missing in the *let*-*381*(*ns1023*) animal (bottom). (G) Quantification of percentage of animals with HMC morphology defects as assessed with the *arg*-*1*promoter::*gfp* reporter. Number of animals (n) scored for each genotype for each reporter is shown under or next to reporter transgene name in (B), (C), (E) and (G). Unpaired *t* test was used for statistical analyses in (C) and (E). n.s. = not significant. a.u. = arbitrary units. Anterior is left, dorsal is up, and scale bars are 10 μm for all animal images.

To assess whether LET-381 is required to maintain HMC morphology, we took advantage of our finding that HMC expression of *arg-1prom::gfp* is not affected in *let- 381(ns1023)* animals (Fig. 3B, E), allowing us to visualize HMC cell shape. We found that 92% of *arg-1prom::gfp; let-381(ns1023)* animals have HMC cell shape defects (Fig. 3F, G). Thus, LET-381 is continuously required in the HMC to maintain gene expression and cell morphology.

### *dmd-4/Dmrt* is also required for HMC fate and regulates HMC gene expression

Our previous studies demonstrated that LET-381 acts in conjunction with one of its targets, the transcription factor UNC-30/Pitx2, to specify GLR glia fate (Stefanakis et al., 2024). Similarly, another study revealed a LET-381-regulated transcription factor, CEH- 34/Six2, acts with LET-381 to drive post-embryonic coelomocyte fate acquisition (Amin et al., 2010). We wondered, therefore, whether a similar regulatory scheme drives HMC development. The Doublesex/Mab3-related transcription factor DMD-4/Dmrt is required for gene expression in the HMC (Bayer et al., 2020, Stefanakis et al., 2024) and its own expression is regulated in the HMC by LET-381 (Fig. 3C, D). Thus, like UNC-30 and CEH-34 in GLR glia and coelomocytes, respectively, DMD-4 might function together with LET-381 for HMC fate specification. To test this idea, we examined expression of five additional HMC gene reporters in *dmd-4*(*ot933*) mutants, lacking sequences encoding the DMD-4 DNA binding domain. While most *dmd-4*(*ot933*) mutants die as embryos, some escape lethality and reach adulthood (Bayer et al., 2020). We found that none of the five HMC reporters we tested, including *let-381::gfp*, is expressed in these escapers (Fig. 4A, B). Furthermore, as in *let-381* mutants, the HMC cell body is also not detected by DIC microscopy in young adult *dmd-4*(*ot933*) mutants (Fig. 4C). These results suggest that the HMC is not specified in the absence of *dmd-4*. Indeed, *dmd-4*(*ot933*) animals exhibit accumulation of food material in their anterior intestine, consistent with a previously-described role for the HMC in controlling the *C. elegans* defecation cycle (Choi et al., 2023).

**FIGURE 4.**
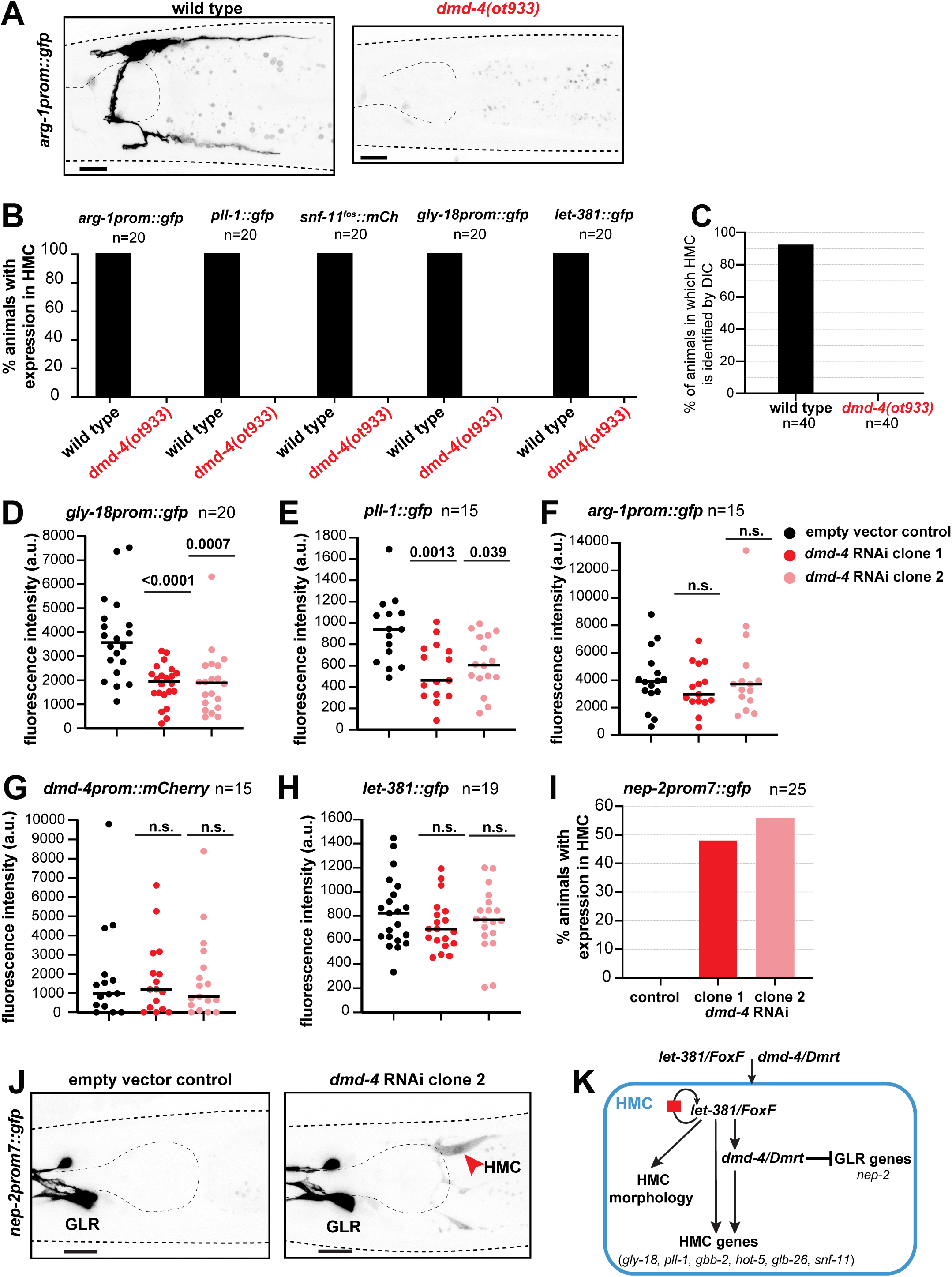
DMD-4 is required for HMC fate specification, HMC gene expression, and repression of GLR glia genes in the HMC. (A) HMC-specific *arg*-*1prom*::*gfp* reporter expression in wild-type (left) and *dmd*-*4*(*ot933*) animals (right). Expression is not observed in *dmd*-*4*(*ot933*). (B) Quantification of percentage of animals with expression in HMC of five different reporters (indicated above the bars) for wild type and *dmd*-*4*(*ot933*). Expression in the HMC is not detected in the *ot933* mutants for all five reporters. (C) Percentage of young adult animals in which the HMC cell body can be identified by DIC microscopy in wild type and *dmd-4(ot933)* mutants. (D) – (H) Quantification of fluorescence intensity for *gly*-*18prom*::*gfp* (D), *pll*- *1*::*gfp* (E), *arg*-*1prom*::*gfp* (F), *dmd*-*4prom*::*mCherry* (G) and *let*-*381*::*gfp* (H) reporters in control and *dmd-4* RNAi animals. (I) Quantification of animals with ectopic HMC expression of the GLR-specific reporter *nep*-*2prom7*::*gfp* upon *dmd-4* RNAi. (J) Representative images showing expression of *nep*-*2prom7*::*gfp* in control and *dmd*- *4(RNAi*) animals. Red arrowhead, HMC. (K) Schematic representation summarizing the regulatory network for HMC development identified in this study. Number of animals (n) scored for each genotype for each reporter is shown under or next to reporter transgene name in (B), (D) – (I) and under the genotype for (C). Unpaired *t* test was used for statistical analysis in (D) - (H). n.s. = not significant. a.u. = arbitrary units. Anterior is left, dorsal is up, and scale bars are 10 μm for all animal images.

To distinguish between fate specification and maintenance roles of DMD-4, we examined the expression of HMC reporters in animals grown postembryonically for three days on bacteria expressing dsRNA against *dmd-4*. We found that induction of RNAi using two different bacterial dsRNA vectors, targeting either a portion of or the entire *dmd- 4* mRNA (*dmd-4* RNAi clone 1 and clone 2, respectively), downregulates expression of the *gly-18prom::gfp* and *pll-1::gfp* HMC reporters, but not of *arg-1prom::gfp* (Fig. 4D-F). Thus, DMD-4 is required for maintaining expression of some HMC genes. Expression of a *dmd-4prom::mCherry* transcriptional reporter is not affected by *dmd-4*(RNAi), suggesting that DMD-4 is not required to maintain its own expression in HMC (Fig. 4G).

Importantly, *let-381::gfp* expression, which is affected in *dmd-4*(*ot933*) mutants, is not affected by *dmd-4*(RNAi), suggesting that as with UNC-30 and CEH-34, DMD-4 acts downstream of LET-381 (Fig. 4H). We conclude, therefore, that DMD-4 acts together with LET-381 in HMC fate specification and maintenance.

### DMD-4 represses expression of some GLR glia genes in the HMC

We previously showed that UNC-30, which acts together with LET-381 in GLR glia, represses expression of HMC genes in these glia (Stefanakis et al., 2024). We wondered, therefore, whether DMD-4 reciprocally represses GLR glia gene expression in the HMC. Indeed, we found that ∼50% of *dmd-4*(RNAi) animals express the GLR glia-specific marker *nep-2prom7::gfp* ectopically in the HMC cell (Fig. 4I, J). This ectopic expression appears dimmer than *nep-2prom7::gfp* expression in GLR glia (Fig. 4J). Four other GLR glia reporters we tested, whose GLR glia expression is lower than *nep-2prom7::gfp*, do not show ectopic expression in the HMC (Fig. S1A), perhaps because of this lower baseline expression. Together, these results suggest that DMD-4 acts to repress at least some GLR glia genes in the HMC.

## DISCUSSION

We describe a gene regulatory network governing fate specification and maintenance of the *C. elegans* endothelial-like HMC (Fig. 4K). Early in development, the LET-381/FoxF transcription factor specifies HMC fate, while later it is continuously required to maintain HMC gene expression and morphology. An autoregulatory sequences upstream of the *let-381* locus ensures continual LET-381 expression. DMD-4/Dmrt, another transcriptional regulator and a target of LET-381, acts with LET-381/FoxF for both HMC fate specification and maintenance. DMD-4, in turn, also represses GLR glia gene expression in the HMC.

*C. elegans* LET-381/FoxF is expressed in only three cell types: GLR glia, which approximate the inner aspect of the central neuropil, the nerve ring, and exhibit astrocyte and endothelial characteristics (Stefanakis et al., 2024); coelomocytes, liver-like detoxifying cells residing within the coelomic cavity (Fares and Greenwald, 2001); and the endothelial-like HMC. All three cell types are non-contractile cells that derive from the mesoderm-like lineage of the MS blast cell. This study, together with previously published results, shows that LET-381/FoxF acts as a master regulator that specifies and maintains all three cell types, and does so by co-regulating a wide variety of cell-type specific genes and cell morphologies (Stefanakis et al., 2024, Amin et al., 2010). How does a single regulator specify such different cell types? Our findings suggest that specificity is enabled via collaboration with co-functional target transcription factors. Specifically, DMD-4/Dmrt, UNC-30/Pitx2, and CEH-34/Six2 act with LET-381/FoxF to control development of the HMC, GLR glia, and coelomocytes, respectively. A similar regulatory strategy is observed in the nervous system, where master regulatory factors, termed Terminal Selectors, combine with different co-acting transcriptional regulators to specify and maintain a variety of neuronal features, including gene expression and connectivity (Allan and Thor, 2015, Hobert and Kratsios, 2019). Indeed, assigning different co-acting transcription factors to a common core transcription factor to form core regulatory complexes (CoRCs), which direct expression of distinct effector gene sets, is a widespread regulatory strategy that can be effectively used to generate novel cell types during evolution (Arendt et al., 2016).

Terminal Selectors typically co-activate many cell-type-specific target genes. Recent findings suggest that apart from gene activation, Terminal Selectors also repress gene expression of alternative cell types (Feng et al., 2020, Remesal et al., 2020). Our findings suggest that such repression is mediated not by the CoRC core-factor, LET-381 here, but by its cofactors, DMD-4 and UNC-30, each repressing gene expression of the sister cell fate instructed by the other. It is thus possible that apart from establishing cell-type specificity, Terminal Selector cofactors safe-guard cell fate through repression of alternative fates.

Terminal Selectors have also been implicated in controlling cell morphology, and our studies here are consistent with this idea, as *let-381* mutants exhibit alterations in HMC cell shape (Fig. 3F). Identification of specific morphology-related HMC target genes could, therefore, unveil molecular mechanisms controlling cell shape.

While Foxf genes have conserved roles in visceral muscle development (Ormestad et al., 2006, Zaffran et al., 2001), roles for these genes in the specification of non-contractile cells are now also emerging. *foxf1* knockdown in *Planaria* results in loss of several cell types, including phagocytic glia and pigment cells (Scimone et al., 2018). In mammals, *Foxf2* is required for differentiation of mesodermal pericytes (Reyahi et al., 2015). Inactivation of murine *Foxf1* results in complete absence of vasculogenesis in the the yolk sac (Mahlapuu et al., 2001). Although this defect appears to be due to an early role of *Foxf1* in splanchnic mesoderm and prior to endothelial cell specification (Astorga and Carlsson, 2007), single cell RNA-seq studies show that *Foxf* genes are enriched in certain endothelial cells in the adult brain and lungs (Hupe et al., 2017, Paik et al., 2020, Wang et al., 2024). Functional roles for these genes in the endothelium have not been investigated. We believe it is possible, and perhaps likely, that as in *C. elegans*, Foxf factors cell autonomously specify and maintain mammalian endothelial cell fates and work together with specific CoRC cofactors to give rise to endothelial cell heterogeneity in different tissues. Indeed, Foxf2, expressed specifically in brain endothelial cells, is sufficient to induce expression of blood-brain barrier associated markers in cell culture (Hupe et al., 2017). However, further investigation is required to clarify whether *Foxf* genes have such roles in vivo.

## MATERIALS AND METHODS

### Caenorhabditis elegans strains and handling

Animals were grown on nematode growth media (NGM) plates seeded with *E. coli* (OP50) bacteria as a food source unless otherwise mentioned. Strains were maintained by standard methods (Brenner, 1974). Wild type is strain N2, *C. elegans* variety Bristol RRID:WB-STRAIN:WBStrain00000001. A complete list of strains generated and used in this study is listed below. A few of the strains were previously published, and/or obtained from the Caenorhabditis Genetic Center (CGC).

### CRISPR/Cas9 genome editing

CRISPR/Cas9 genome editing was performed using Cas9, tracrRNAs and crRNAs from IDT as described in (Dokshin et al., 2018). Generation of the *let-381*(*ns1023*) and the previously described *let*-*381*(*ns1026*) deletion alleles was performed by use of two crRNAs (tggttgaagagacatacatc, ttatggatggaaaacagacg) and a ssODN (tcatcatacttttccctctatcttctcaaccagatctgttttccatccataagccaccaccccattctgc). CRISPR/Cas9 generated different deletion alleles: *ns1023* carrying a 107 bp deletion from –415 to –308 and *ns1026,* an indel carrying a deletion from –481 to –340 and insertion of a random 34 bp sequence cttatcttctcaatcttctcaaccagatgtgttg. Both alleles remove the tgtttata *let-381* motif at -333 bp from the ATG.

### Generation of transgenic reporters

The *glb*-*26prom1*::*gfp* reporter was generated by a PCR fusion approach (Hobert, 2002). Genomic promoter fragments were fused to *gfp* followed by the *unc-54* 3’ UTR. Promoters were initially amplified with primers A (gactgtggagacgatcgtac) and B (CTCTAGAGTCGACCTGCAGGCATGCAAGCTctgggaatgagcacacgaaa) from N2 genomic DNA and *gfp* followed by *unc-54* 3’UTR was amplified by primers C (AGCTTGCATGCCTGCAGGTCG) and D (AAGGGCCCGTACGGCCGACTA) from plasmid pPD9575. For the fusion step, PCR amplification was performed using primers A* (gttcgaagatctgcacgaag) and D* (CAAGAAAAACGCCGTCCTCG) as previously described (Hobert, 2002). PCR fusion DNA fragments were injected as simple extrachromosomal arrays in the wildtype N2 strain in the following concentrations: *glb*- *26prom1*::*gfp,* 50ng/μL, *myo-3prom::mCherry* (co-injection marker), 25ng/μL, pBluescript SK+, 25ng/μL.

### Generation of new *dmd-4* RNAi clone

A 4544 bp fragment containing the entire *dmd*-*4* genomic locus was PCR amplified from N2 genomic DNA with primers AGACCGGCAGATCTGATATCATCGATGAATTCGAGCTCCAgtcgagctccgcctacaatc and GCGCGTAATACGACTCACTATAGGGCGAATTGGGTACCGGgaggggatttgccacaagta and subsequently cloned by Gibson cloning into the empty RNAi vector L4440, which was PCR amplified by primers CCGGTACCCAATTCGCCCTA and TGGAGCTCGAATTCATCGAT. Ligated plasmids were then transformed into HT115 *E. coli* bacteria by electroporation. Plasmid clones containing the properly inserted *dmd-4* locus were identified by Whole Plasmid sequencing, performed by Plasmidsaurus using Oxford Nanopore Technology with custom analysis and annotation.

### RNAi by feeding

Synchronized L1 larvae were placed on NGM plates with 1mM IPTG and 25μg/mL carbenicillin, and coated with bacteria carrying either the *dmd*-*4* RNAi or the empty vector RNAi control plasmids. Worms were grown for three days at 20°C and then mounted on agarose slides for imaging on a compound microscope as described in the Microscopy section below. HMC is refractory to RNAi; thus, RNAi sensitized background strains carrying *eri-1(mg366)* (Kennedy et al., 2004) were used for these experiments.

### Microscopy

Animals were anesthetized using 100mM NaN3 (sodium azide) and mounted on 5% agarose pads on glass slides. Z-stack images (each ∼7μm thick) were acquired using either a Zeiss confocal microscope LSM990 (images in Figs. 1B, 1F, 2B, 3F, 4A, 4J) or a Zeiss compound microscope Axio Imager M2 (images in Figs. 1D, 3A, 3D) using MicroManager software (version 1.4.22) (Edelstein et al., 2010). ImageJ (Schneider et al., 2012) was used to produce maximum projections of z-stack images (2–20 slices) presented in the Figures. Figures were prepared by using Adobe Illustrator.

### Quantification and Statistical Analysis

All microscopy fluorescence quantifications were done in ImageJ (Schneider et al., 2012). For each experiment, Mutant (or RNAi) and control animals were imaged during the same imaging session with all acquisition parameters maintained constant between the two groups. Fluorescence intensity of gene expression in the HMC cell (Figs. 3C, 3E, 4D – H) was measured in the plane with strongest signal within the z-stack in a region drawn around the HMC nucleus (for nuclear reporters) or cell body (for cytoplasmic reporters). A single circular region in an adjacent area was used to measure background intensity for each animal; this value was then subtracted from the fluorescence intensity of reporter expression for each HMC cell. Quantification of percentage of animals with reporter expression in HMC (Figs. 1C, 2C, 3B, 4B, 4I, S1A), quantification of percentage of animals in which HMC can be identified by Differential Interference Contrast (DIC) microscopy (1E, 4C) and quantification of percentage of HMC with morphology defects (Fig. 3G) were performed by manual counting using ImageJ. Prism (GraphPad) was used for graphs and statistical analysis as described in Figure Legends. Unpaired two-sided Students’ t test was used to determine the statistical significance between two groups.

## List of strains used and generated in this study

| Strain Name | Genotype | Reference |
| --- | --- | --- |
| N2 | wild type | Caenorhabditic Genetics Center (CGC) |
| VC706 | *let-381(gk302) I / hT2 (I;III)* | (Barstead et al., 2012) |
| KR429 | *dpy-5(e61) let-381(h107) unc-13(e450) I ; sDp2 (I:f)* | (Howell et al., 1987) |
| GR1373 | *eri-1(mg366) IV* | (Kennedy et al., 2004) |
| DE60 | *dnIs13 [gly-18prom::gfp + unc-119(+)] I ; unc-119(e2498) III* | (Warren et al., 2001) |
| MT20492 | *lin-15B&lin-15A(n765) X ; nIs471 [lgc-55prom::gfp + lin-15(+)]* | (Ringstad et al., 2009) |
| OH13027 | *otIs569 [snf-11(fosmid)::SL2::H2B::mChopti + pha-1(+)]* | (Gendrel et al., 2016) |
| PD4443 | *ccls4443 [arg-1prom::gfp + dpy-20(+)] IV* | (Kostas and Fire, 2002) |
| SD1633 | *ccls4251 [(pSAK2) myo-3p::GFP::LacZ::NLS + (pSAK4) myo-3p::mitochondrial GFP + dpy-20(+)] I ; stIs10539 [dmd-4p::HIS-24::mCherry + unc-119(+)]* | (Liu et al., 2009) |
| OH16286 | *dmd-4(ot933) X* | (Bayer et al., 2020) |
| OS14679 | *nsEx7257 [glb-26prom1::gfp + myo-3::mCherry] line 1* | This study |
| OS14681 | *nsEx7258 [glb-26prom1::gfp + myo-3::mCherry] line 2* | This study |
| OS14683 | *nsEx7259 [glb-26prom1::gfp + myo-3::mCherry] line 3* | This Study |
| OS11703 | *nsIs746 [nep-2prom7::gfp] V* | (Stefanakis et al., 2024) |
| OS12099 | *nsIs831 [pll-1prom1::tagrfp] X* | (Stefanakis et al., 2024) |
| OS12103 | *nsIs835 [hot-5prom2::gfp] X* | (Stefanakis et al., 2024) |
| OS12700 | *unc-30(ns959[unc-30::gfp::degron(AID)]) IV* | (Stefanakis et al., 2024) |
| OS13288 | *let-381(ns995[let-381::gfp::aid]) I* | (Stefanakis et al., 2024) |
| PHX3025 | *hlh-1(syb3025[hlh-1::gfp::aid]) II* | (Stefanakis et al., 2024) |
| PHX5792 | *pll-1(syb5792[pll-1::sl2::gfp::h2b]) III* | (Stefanakis et al., 2024) |
| PHX5759 | *gbb-2(syb5759[gbb-2::sl2::gfp::h2b]) IV* | (Stefanakis et al., 2024) |
| OS13835 | *let-381(ns1023[\*ns995]) I* | This study |
| OS13838 | *let-381(ns1026[\*ns995]) I* | (Stefanakis et al., 2024) |

## AUTHOR CONTRIBUTIONS

**Nikolaos Stefanakis**: Conceptualization; Methodology; Formal Analysis; Investigation; Writing – Original Draft; Writing – Review and Editing; Visualization; Funding Acquisition. **Jasmine Xi**: Investigation; Formal Analysis **Jessica Jiang**: Investigation Formal Analysis; **Shai Shaham**: Conceptualization; Writing – Original draft; Writing – Review and Editing; Project Administration; Funding Acquisition.

## ACKNOWLEDGEMENTS

We thank Oliver Hobert for strains; Members of the Shaham lab for experimental advice, comments and discussion. Some strains were provided by the CGC, which is funded by NIH Office of Research Infrastructure Programs (P40 OD010440).

## DATA AVAILABILITY

- This paper does not report original code.
- All newly generated critical strains will be available at the Caenorhabditis Genetics Center (CGC).
- All Strains and plasmids are available upon request. Any additional information required to reanalyze the data reported and requests for resources and reagents should be directed to and will be fulfilled by the corresponding author.

---

**Supplemental Figure S1.**
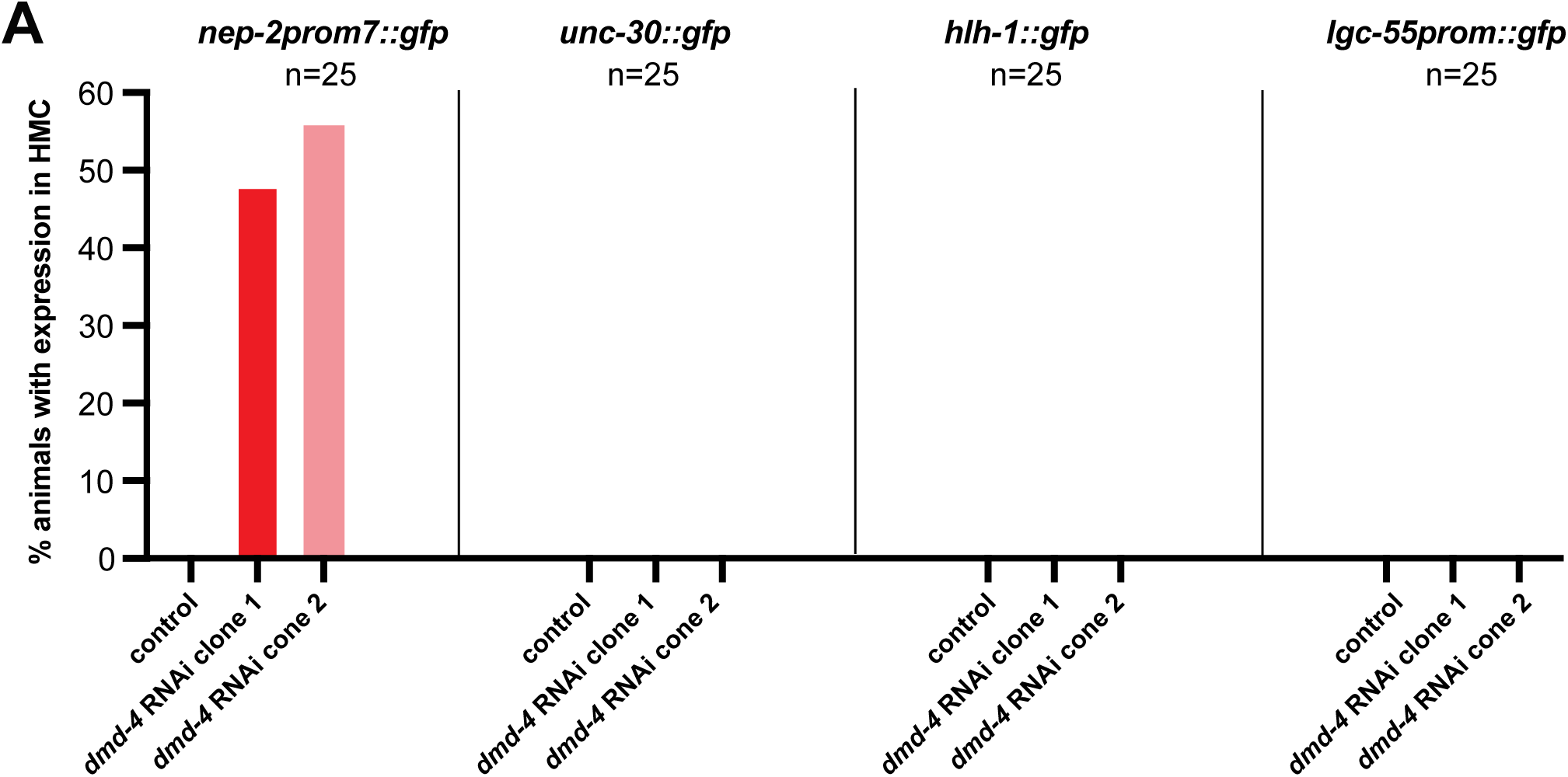
DMD-4 suppresses GLR glia gene expression in the HMC. (A) Quantification of animals with ectopic HMC expression of four GLR glia reporters. Ectopic expression is observed only for *nep*-*2prom7*::*gfp* but not for the other reporters tested. Number of animals (n) tested for each genotype for each reporter is shown under the name of each reporter transgene.

## REFERENCES

1. Allan, D. W. & Thor, S. 2015. Transcriptional selectors, masters, and combinatorial codes: regulatory principles of neural subtype specification. WIREs Developmental Biology, 4, 505–528.

2. Ambros, V. 2011. MicroRNAs and developmental timing. Current Opinion in Genetics & Development, 21, 511–517.

3. Amin, N. M., Shi, H. & Liu, J. 2010. The FoxF/FoxC factor LET-381 directly regulates both cell fate specification and cell differentiation in C. elegans mesoderm development. Development, 137, 1451–60.

4. Arendt, D., Musser, J. M., Baker, C. V. H., Bergman, A., Cepko, C., Erwin, D. H., Pavlicev, M., Schlosser, G., Widder, S., Laubichler, M. D. & Wagner, G. P. 2016. The origin and evolution of cell types. Nature Reviews Genetics, 17, 744–757.

5. Astorga, J. & Carlsson, P. 2007. Hedgehog induction of murine vasculogenesis is mediated by Foxf1 and Bmp4. Development, 134, 3753–3761.

6. Barstead, R., Moulder, G., Cobb, B., Frazee, S., Henthorn, D., Holmes, J., Jerebie, D., Landsdale, M., Osborn, J., CHERILYN Pritchett, Robertson, J., Rummage, J., Stokes, E., Vishwanathan, M., Mitani, S., Gengyo-Ando, K., Funatsu, O., Hori, S., Imae, R., Kage-Nakadai, E., Hiroyuki Kobuna, Machiyama, E., Motohashi, T., Otori, M., Suehiro, Y., Yoshina, S., Smith, M., Moerman, D., Edgley, M., Adair, R., Allan, B. J., Au, V., Chaudhry, I., Cheung, R., Dadivas, O., Eng, S., Fernando, L., Fisher, A., Flibotte, S., Gilchrist, E., Hay, A., Huang, P., Hunt, R. W., Kwitkowski, C., Lau, J., Lee, N., Liu, L., Lorch, A., Luck, C., Maydan, J., Mckay, S., Miller, A., Mullen, G., Navaroli, C., Neil, S., Hunt-Newbury, R., Partridge, M., Perkins, J., Rankin, A., Raymant, G., Rezania, N., Rogula, A., Shen, B., Stegeman, G., Tardif, A., Taylor, J., Veiga, M., Wang, T. & Zapf, R. 2012. Large-scale screening for targeted knockouts in the Caenorhabditis elegans genome. G3 (Bethesda), 2, 1415–25.

7. Bayer, E. A., Stecky, R. C., Neal, L., Katsamba, P. S., Ahlsen, G., Balaji, V., Hoppe, T., Shapiro, L., Oren-Suissa, M. & Hobert, O. 2020. Ubiquitin-dependent regulation of a conserved DMRT protein controls sexually dimorphic synaptic connectivity and behavior. Elife, 9.

8. Brenner, S. 1974. The genetics of Caenorhabditis elegans. Genetics, 77, 71–94.

9. Choi, U., Hu, M., Zhang, Q. & Sieburth, D. 2023. The head mesodermal cell couples FMRFamide neuropeptide signaling with rhythmic muscle contraction in C. elegans. Nature Communications, 14, 4218.

10. Dokshin, G. A., Ghanta, K. S., Piscopo, K. M. & Mello, C. C. 2018. Robust Genome Editing with Short Single-Stranded and Long, Partially Single-Stranded DNA Donors in Caenorhabditis elegans. Genetics, 210, 781–787.

11. Edelstein, A., Amodaj, N., Hoover, K., Vale, R. & Stuurman, N. 2010. Computer control of microscopes using µManager. Curr Protoc Mol Biol, Chapter 14, Unit14.20.

12. Fares, H. & Greenwald, I. 2001. Genetic Analysis of Endocytosis in Caenorhabditis elegans: Coelomocyte Uptake Defective Mutants. Genetics, 159, 133–145.

13. Feng, W., Li, Y., Dao, P., Aburas, J., Islam, P., Elbaz, B., Kolarzyk, A., Brown, A. E. & Kratsios, P. 2020. A terminal selector prevents a Hox transcriptional switch to safeguard motor neuron identity throughout life. Elife, 9.

14. Figueroa, X. F. & Duling, B. R. 2009. Gap junctions in the control of vascular function. Antioxid Redox Signal, 11, 251–66.

15. Gendrel, M., Atlas, E. G. & Hobert, O. 2016. A cellular and regulatory map of the GABAergic nervous system of C. elegans. Elife, 5.

16. Gieseler, K., Qadota, H. & Benian, G. M. 2017. Development, structure, and maintenance of C. elegans body wall muscle. WormBook, 2017, 1–59.

17. Griffith, T. M. 2004. Endothelium-dependent smooth muscle hyperpolarization: do gap junctions provide a unifying hypothesis? Br J Pharmacol, 141, 881–903.

18. Hall, D. H. & Altun, Z. F. 2008. C. Elegans Atlas, Cold Spring Harbor Laboratory Press.

19. Hennigs, J. K., Matuszcak, C., Trepel, M. & Körbelin, J. 2021. Vascular Endothelial Cells: Heterogeneity and Targeting Approaches. Cells, 10, 2712.

20. Hobert, O. 2002. PCR fusion-based approach to create reporter gene constructs for expression analysis in transgenic C. elegans. Biotechniques, 32, 728–30.

21. Hobert, O. & Kratsios, P. 2019. Neuronal identity control by terminal selectors in worms, flies, and chordates. Curr Opin Neurobiol, 56, 97–105.

22. Horowitz, L. B. & Shaham, S. 2024. Apoptotic and Nonapoptotic Cell Death in Caenorhabditis elegans Development. Annual Review of Genetics, 58, 113–134.

23. Howell, A. M., Gilmour, S. G., Mancebo, R. A. & Rose, A. M. 1987. Genetic analysis of a large autosomal region in Caenorhabditis elegans by the use of a free duplication. Genetics Research, 49, 207–213.

24. Hupe, M., Li, M. X., Kneitz, S., Davydova, D., Yokota, C., Kele, J., Hot, B., Stenman, J. M. & Gessler, M. 2017. Gene expression profiles of brain endothelial cells during embryonic development at bulk and single-cell levels. Sci Signal, 10.

25. Kennedy, S., Wang, D. & Ruvkun, G. 2004. A conserved siRNA-degrading RNase negatively regulates RNA interference in C. elegans. Nature, 427, 645–9.

26. Kostas, S. A. & Fire, A. 2002. The T-box factor MLS-1 acts as a molecular switch during specification of nonstriated muscle in C. elegans. Genes Dev, 16, 257–69.

27. Kratsios, P. & Hobert, O. 2024. Almost 40 years of studying homeobox genes in C. elegans. Development, 151.

28. Liu, X., Long, F., Peng, H., Aerni, S. J., Jiang, M., Sánchez-Blanco, A., Murray, J. I., Preston, E., Mericle, B., Batzoglou, S., Myers, E. W. & Kim, S. K. 2009. Analysis of Cell Fate from Single-Cell Gene Expression Profiles in C. elegans. Cell, 139, 623–633.

29. Maguire, J. J. & Davenport, A. P. 2005. Regulation of vascular reactivity by established and emerging GPCRs. Trends in Pharmacological Sciences, 26, 448–454.

30. Mahlapuu, M., Ormestad, M., Enerback, S. & Carlsson, P. 2001. The forkhead transcription factor Foxf1 is required for differentiation of extra- embryonic and lateral plate mesoderm. Development, 128, 155–66.

31. Marziano, C., Genet, G. & Hirschi, K. K. 2021. Vascular endothelial cell specification in health and disease. Angiogenesis, 24, 213–236.

32. Olsson, A.-K., Dimberg, A., Kreuger, J. & Claesson-Welsh, L. 2006. VEGF receptor signallingin control of vascular function. Nature Reviews Molecular Cell Biology, 7, 359–371.

33. Ormestad, M., Astorga, J., Landgren, H., Wang, T., Johansson, B. R., Miura, N. & Carlsson, P. 2006. Foxf1 and Foxf2 control murine gut development by limiting mesenchymal Wnt signaling and promoting extracellular matrix production. Development, 133, 833–43.

34. Paik, D. T., Tian, L., Williams, I. M., Rhee, S., Zhang, H., Liu, C., Mishra, R., Wu, S. M., Red-Horse, K. & Wu, J. C. 2020. Single-Cell RNA Sequencing Unveils Unique Transcriptomic Signatures of Organ-Specific Endothelial Cells. Circulation, 142, 1848–1862.

35. Phng, L. K. & Gerhardt, H. 2009. Angiogenesis: A Team Effort Coordinated by Notch. Developmental Cell, 16, 196–208.

36. Remesal, L., Roger-Baynat, I., Chirivella, L., Maicas, M., Brocal-Ruiz, R., Perez-Villalba, A., Cucarella, C., Casado, M. & Flames, N. 2020. PBX1 acts as terminal selector for olfactory bulb dopaminergic neurons. Development, 147.

37. Reyahi, A., Nik, A. M., Ghiami, M., Gritli-Linde, A., Ponten, F., Johansson, B. R. & Carlsson, P. 2015. Foxf2 Is Required for Brain Pericyte Differentiation and Development and Maintenance of the Blood-Brain Barrier. Dev Cell, 34, 19–32.

38. Richmond, J. 2005. Synaptic function. WormBook, 1–14.

39. Ringstad, N., Abe, N. & Horvitz, H. R. 2009. Ligand-gated chloride channels are receptors for biogenic amines in C. elegans. Science, 325, 96–100.

40. Rohlenova, K., Veys, K., Miranda-Santos, I., de Bock, K. & Carmeliet, P. 2018. Endothelial Cell Metabolism in Health and Disease. Trends in Cell Biology, 28, 224–236.

41. Schneider, C. A., Rasband, W. S. & Eliceiri, K. W. 2012. NIH Image to ImageJ: 25 years of image analysis. Nature Methods, 9, 671–675.

42. Scimone, M. L., Wurtzel, O., Malecek, K., Fincher, C. T., Oderberg, I. M., Kravarik, K. M. & Reddien, P. W. 2018. foxF-1 Controls Specification of Non-body Wall Muscle and Phagocytic Cells in Planarians. Curr Biol, 28, 3787–3801 e6.

43. Singhvi, A., Shaham, S. & Rapti, G. 2024. Glia Development and Function in the Nematode Caenorhabditis elegans. Cold Spring Harb Perspect Biol.

44. Stefanakis, N., Jiang, J., Liang, Y. & Shaham, S. 2024. LET-381/FoxF and its target UNC-30/Pitx2 specify and maintain the molecular identity of *C. elegans* mesodermal glia that regulate motor behavior. The EMBO Journal, 43, 956–992.

45. Trimm, E. & Red-Horse, K. 2023. Vascular endothelial cell development and diversity. Nature Reviews Cardiology, 20, 197–210.

46. Wang, G., Wen, B., Guo, M., Li, E., Zhang, Y., Whitsett, J. A., Kalin, T. V. & Kalinichenko, V. V. 2024. Identification of endothelial and mesenchymal FOXF1 enhancers involved in alveolar capillary dysplasia. Nature Communications, 15, 5233.

47. Warren, C. E., Krizus, A. & Dennis, J. W. 2001. Complementary expression patterns of six nonessential Caenorhabditis elegans core 2/I N- acetylglucosaminyltransferase homologues. Glycobiology, 11, 979–988.

48. Zaffran, S., Kuchler, A., Lee, H. H. & Frasch, M. 2001. biniou (FoxF), a central component in a regulatory network controlling visceral mesoderm development and midgut morphogenesis in Drosophila. Genes Dev, 15, 2900–15.

